# Uncovering Latent Biological Function Associations through Gene Set Embeddings

**DOI:** 10.1101/2024.10.10.617577

**Authors:** Yuhang Huang, Fan Zhong

**Affiliations:** Institutes of Biomedical Sciences, Fudan University, 131 Dongan Road, Shanghai, Shanghai 200032, China; Intelligent Medicine Institute, Fudan University, Shanghai 200032, China

**Keywords:** Biological network analysis, MSigDB, Gene-term associations, Cross-species analysis, Network embedding

## Abstract

The complexity of biological systems has increasingly been unraveled through computational methods, with biological network analysis now focusing on the construction and exploration of well-defined interaction networks. Traditional graph-theoretical approaches have been instrumental in mapping key biological processes using high-confidence interaction data. However, these methods often struggle with incomplete or/and heterogeneous datasets. In this study, we extend beyond conventional bipartite models by integrating attribute-driven knowledge from the Molecular Signatures Database (MSigDB) using the node2vec algorithm. Our approach explores unsupervised biological relationships and uncovers potential associations between genes and biological terms through network connectivity analysis. By embedding both human and mouse data into a shared vector space, we validate our findings cross-species, further strengthening the robustness of our method. This integrative framework reveals both expected and novel biological insights, offering a comprehensive perspective that complements traditional biological network analysis and paves the way for deeper understanding of complex biological processes and diseases.

## Introduction

The intricate architecture of biological systems has been progressively demystified through advanced computational methodologies. Currently, biological network analysis predominantly focuses on constructing and interrogating well-defined interaction networks. These approaches often employ graph-theoretical methods to identify key nodes and critical pathways, utilizing high-confidence interaction data to map out precise biological processes. Databases such as KEGG (Kanehisa, Furumichi, Tanabe, Sato, & Morishima, 2017) and STRING (Szklarczyk *et al*., 2019) are commonly used.

Initially, researchers employed network-based methods primarily to analyze biological data (Dybowski & Gant, 1995). Subsequent advancements have profoundly deepened our understanding of biological networks through unsupervised analysis. Jeong *et al*. (Jeong, Tombor, Albert, Oltvai, & Barabási, 2000) unveiled complex structures within metabolic networks, Barabási and Oltvai (Barabási & Oltvai, 2004) elucidated the functional organization of cellular networks, and Stuart *et al*. (Stuart, Segal, Koller, & Kim, 2003) were pioneers in identifying conserved gene modules across species.

Cumulatively, these seminal studies underscore the evolution and significant impact of network analysis in bioinformatics. Within the past decades, researchers have been applying machine learning on graphs and utilizing various methods to represent (W. Hamilton, Z. Ying, & J. Leskovec, 2017): from shallow embedding approaches like DeepWalk (Perozzi, Al-Rfou, & Skiena, 2014), node2vec (Grover & Leskovec, 2016), and HARP (Chen, Perozzi, Hu, & Skiena, 2018), to methods based on deep learning like neighborhood autoencoder methods DNGR (Cao, Lu, & Xu, 2016), SDNE (Wang, Cui, & Zhu, 2016), and convolutional one like GraphSAGE (W. L. Hamilton, R. Ying, & J. Leskovec, 2017).

Among these methods, node2vec stands out due to its simplicity and strong interpretability in an unsupervised learning context. In the field of computational biology, graph representation learning has gained popularity in the analysis of biological relationships. Numerous studies have reviewed and summarized the applications of graph representation learning in the biomedical field (Nelson *et al*., 2019; Yi, You, Huang, & Kwoh, 2021), demonstrating its versatility. These approaches typically model biological systems through bipartite (Zhang *et al*., 2020), multirelational (Y. Liu, Zeng, He, & Zou, 2017), or knowledge-based graphs (Rotmensch, Halpern, Tlimat, Horng, & Sontag, 2017), which capture intricate relationships between biological entities, molecules, and biomedical concepts, providing powerful tools for tasks such as drug discovery (Zong *et al*., 2021), disease treatment, and molecular interaction analysis (Fachet *et al*., 2020).

Building on advancements in graph-based representation learning, this study extends traditional bipartite models by incorporating attribute-driven knowledge database, Molecular Signatures Database (MsigDB, Liberzon *et al*., 2015). Traditional biological network analyses, while effective with specific, well-curated datasets, often falter when faced with incomplete or ambiguous data from diverse sources (R. Liu, Hirn, & Krishnan, 2023). To bridge this gap, we introduce a novel network-based framework designed to explore complex relationships within biological terms (Figure 1). Rather than relying solely on annotated data, our approach uncovers potential gene associations with functional terms (e.g., pathways, functions) through network connectivity. By analyzing network structure and topology, we aim to reveal hidden patterns and novel associations, offering deeper insights, particularly in exploratory studies where direct evidence is limited or mechanisms are unclear. This integrative approach enhances traditional data mining by offering a more comprehensive perspective.

**Figure 1.**
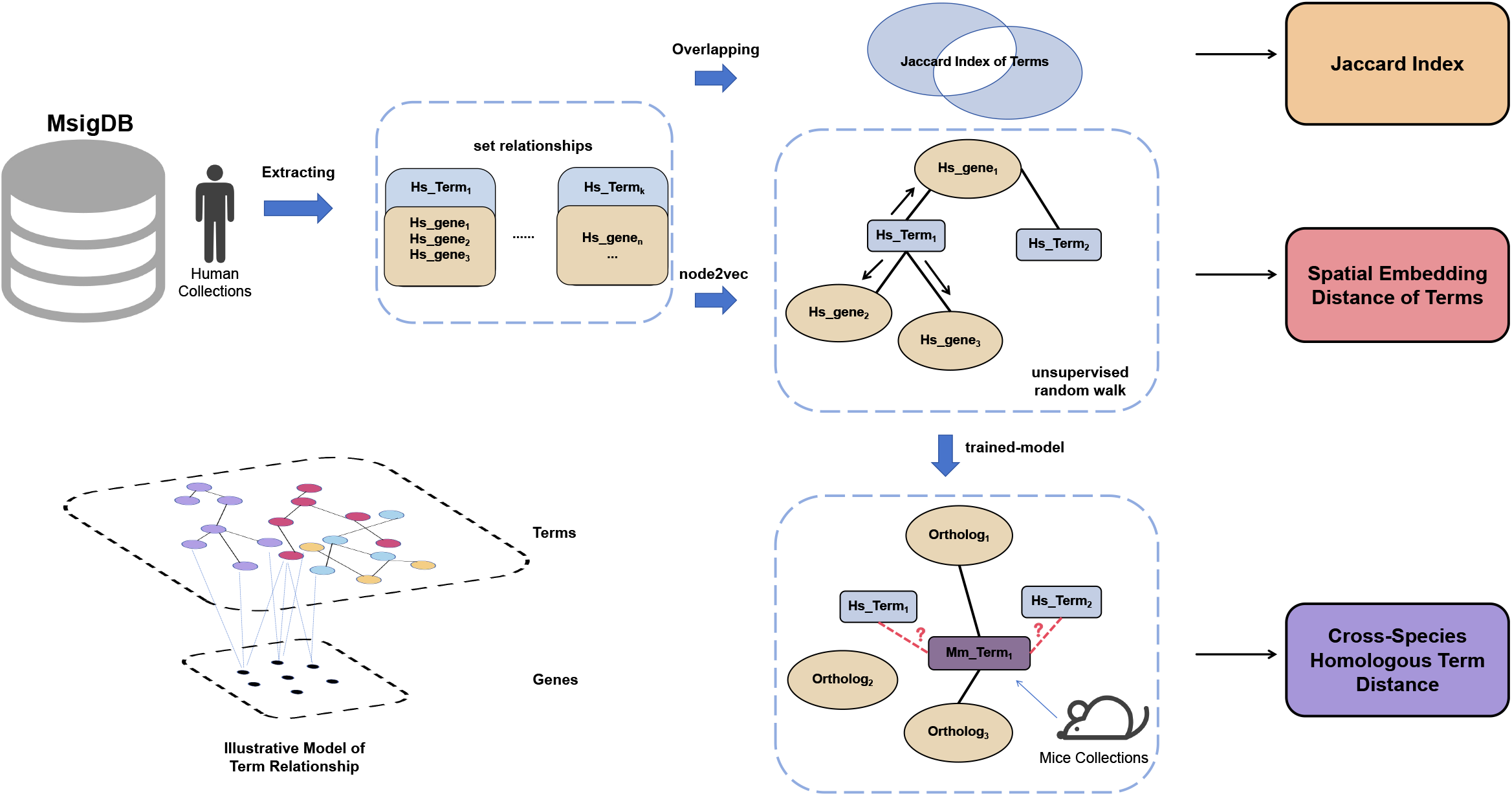
Cross-species validation pipeline: We utilized the Jaccard index and node2vec-based distance for analysis, embedding both human and mouse datasets into a shared vector space. Aligning homologous genes across species revealed consistent patterns and reinforced the robustness of our findings. We focused solely on the relationships between terms, and the diagram in the lower left is an abstract representation of this concept.

## Method

### Data Integration

The dataset incorporates a comprehensive set of terms that encapsulate a wide array of biological phenomena, we have opted to focus on a single species to mitigate these potential confounders and to ensure the integrity of the analysis. Within this scope, we define two essential sets: the set of **genes**, denoted as *G* = { *g*_1_, *g*_2_, . . ., *g*_*m*_ }, and the set of **terms**, denoted as *T* = {*t*_1_,*t*_2_, . . ., *t*_*n*_}. Data for these sets were sourced from the MSigDB (https://www.gsea-msigdb.org/gsea/msigdb). To bridge species-specific differences in gene nomenclature, we utilized the biomaRt (Durinck, Spellman, Birney, & Huber, 2009) R package, which facilitated the conversion of murine genes to their human orthologs, ensuring cross-species comparability in our analysis.

Following data preprocessing, we construct an index dictionary to facilitate the creation of a PyG (Fey & Lenssen, 2019) (PyTorch Geometric) data object. This entails defining two mappings: one from genes to indices, termed gene_to_index: *G* → {0,1, . . ., *m* − 1}, and another from terms to indices, termed term_to_index: *T* → { *m, m* + 1, . . ., *m* + *n* − 1}.

We then generate the list of edges E by adding an edge for each gene-term pair (*g,t*) to form the edge set *E* = {(gene_to_index(*g*), term_to_index(*t*)}. The PyG data object, represented as *Graph*, encompasses the node set *V* = *G* ∪ *T*, and the edge set *E*: *Graph* = (*V, E*), where the number of nodes |*V*| is equal to *m* + *n*.

### Reliability Score

Our analysis revealed that both the *Jaccard index* and node2vec embeddings distances exhibited extreme value distributions .that across 10^8^ pairwise comparisons. To assess significant biological associations, we normalized both metrics and selected representative pairs at distance thresholds of 0.1, 0.3, 0.5, 0.7, and 1 for further manual evaluation and literature-based validation. For each threshold, 10 pairs were analyzed in-depth to determine their biological relevance. To enhance the rigor of biological relevance assessment, we introduced a reliability score (RS) ranging from 0 to 5, reflecting levels of confidence from speculative associations to those strongly supported by extensive literature (Supplementary Material 1).

### Distance Calculation Methodology

We utilized the node2vec algorithm to embed nodes within our graph-structured data, capturing their contextual relationships. The node2vec algorithm learns these embeddings by performing biased random walks on the graph, balancing between breadth-first search (BFS) and depth-first search (DFS) through two parameters: *p* (return parameter) and *q* (in-out parameter).

The random walks are controlled by the following transition probability formula:

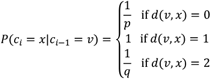

where *c*_*i*_ is the candidate note at step *i, c*_*i* −1_ = *v* is the previous node in the walk, *d*(*v,x*) is the shortest path distance between nodes *v* and *x*,

The *Jaccard index* measures the overlap between two sets, and in our context, it represents the similarity between terms based on the overlap of their corresponding gene sets. Consider two terms, *T*_*A*_ and *T*_*B*_, each associated with a set of genes based on their adjacency in the graph structure.

The *Jaccard Index* between these two terms is calculated using the formula:

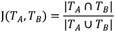

### Node2vec Training and Hyperparameter Optimization

To uncover the underlying patterns within our data, which involve complex many-to-many relationships without precise numerical values, we have adopted an unsupervised learning approach. Central to this approach is the node2vec algorithm, as implemented in the torch_geometric library, which we integrated into a machine learning pipeline. This pipeline enables the sequential execution of node embeddings and subsequent clustering.

Clustering is essential in unsupervised learning as it allows us to organize data points into groups based on their inherent similarities, without relying on pre-labeled classes. The hyperparameter tuning for the node2vec model involved varying several parameters. Embedding dimensions were set at either 32 and 64. Walk lengths were tested at values of 10, 20, and 30. The return parameter *p* and in-out parameter *q* were each explored across the values 0.25, 0.5, 1, 2, and 4. Context size was evaluated at 10 and 20, while the number of negative samples was tested at 1, 5, and 10. Finally, the number of walks per node was adjusted to 10, 20, or 30. This comprehensive hyperparameter search ensured an optimized configuration for the model.

Following the optimization of hyperparameters, we applied *k*-means clustering to evaluate the quality of the embeddings. The cluster labels assigned by *k*-means were used to calculate two internal clustering validation metrics, which assess the coherence and separation of the clusters.

The Silhouette score (Rousseeuw, 1987) ranges from -1 to 1, with higher values indicating better clustering performance. It is defined as:

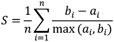

Where *a*_*i*_ is the average distance between the *i*-th data point and other points in the same cluster, *b*_*i*_ is the minimum average distance from the *i* -th data point to the points in a different cluster, *n* is the total number of data points.

The Davies-Bouldin score (Davies & Bouldin, 1979) measures the average similarity between clusters, with lower values indicating better clustering performance. It is calculated by:

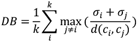

Where *σ*_*i*_ is the average distance of all points in cluster *i* to the cluster centroid, *d*(*c*_*i*_, *c*_*j*_) *i*s the distance between centroids of clusters *i* and *j, k* is the number of clusters.

The 64-dimensional space is more likely to exhibit biological credibility, as confirmed by manual validation and a literature review (Figure 4). Furthermore, we observed that the 32-dimensional embeddings produce more less credible results within the top 5%. Ultimately, we utilized the 64-dimensional embeddings for the subsequent statements of node2vec-based distances.

The model was trained on a server equipped with an NVIDIA GeForce RTX 3080 Ti GPU, using CUDA for parallel computations. The server ran Ubuntu 20.04 LTS and had 256 GB of RAM. We used Python 3.8 and the torch_geometric library for our implementation. The hyperparameter search was conducted over the course of one week, with each parameter combination undergoing 40 epochs of training. The duration of training for each parameter set varied depending on the specific parameters chosen, with some combinations taking more time to complete.

## Results

### Interpreting Biological Relevance of Jaccard Index

A higher *Jaccard index* generally signifies a substantial overlap between gene sets, indicating a robust biological connection. However, lower *Jaccard index* should not be overlooked, particularly when the gene sets involved contain fewer genes, where even modest overlap might indicate meaningful associations. To refine the analysis, we applied a hypergeometric distribution test with a p value threshold of 0.05 to assess the significance of shared gene counts. From the analysis, it is evident that higher *Jaccard index* are generally associated with higher RS (Figure 2), reflecting a trend that aligns with our expectations.

**Figure 2.**
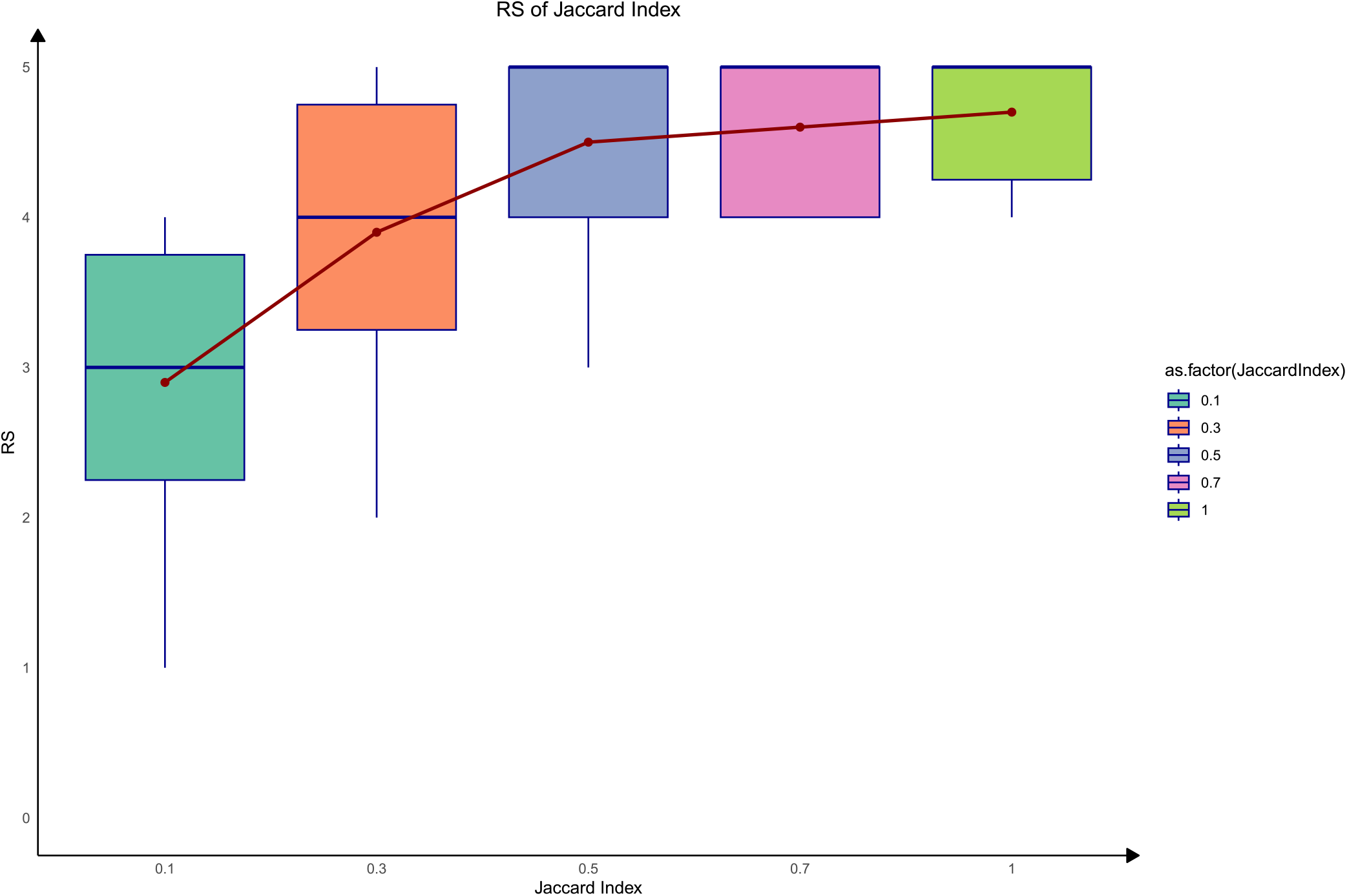
Box plot showing the relationship between the Jaccard index and reliability scores, where higher Jaccard values tend to correspond to higher reliability scores.

For example, the association between GOBP_UBIQUITIN_DEPENDENT_GLYCOPROTEIN_ERAD_PATHWAY and GOMF_MANNOSYL_OLIGOSACCHARIDE_1_2_ALPHA_MANNOSIDASE_ACTIVITY (with a *Jaccard index* of 1) is biologically intuitive. Although this connection is not explicitly indicated in MSigDB, these terms describe closely related processes. The ubiquitin-dependent glycoprotein ERAD pathway is responsible for recognizing and degrading misfolded glycoproteins in the endoplasmic reticulum (ER) via the ER-associated degradation (ERAD) system (Lopata, Kniss, Löhr, Rogov, & Dötsch, 2020). During this process, mannosyl oligosaccharide 1,2-alpha mannosidase enzymes trim misfolded glycoproteins, preparing them for ubiquitination and degradation. The inclusion of this enzyme activity within the pathway underscores the functional cohesion, highlighting the expected biological relationship between the two functional terms.

As the Jaccard values decrease, a turning point emerges around 0.5, where RS typically range between 3 and 4 (Figure 3). Although direct literature support becomes sparse, these associations remain biologically plausible. A notable example is the interaction between FU_INTERACT_WITH_ALKBH8 and GOCC_CHAPERONIN_CONTAINING_T_COMPLEX (CCT). While no explicit link has been documented between the CCT complex and ALKBH8, both share a key gene, *TCP1*, suggesting a potential functional connection. CCT, a type II molecular chaperone in eukaryotes, plays a pivotal role in folding nascent proteins, particularly during T-cell activation and in response to external stressors. Its critical function in maintaining protein homeostasis means that CCT deficiency can lead to immune dysfunction and impaired stress response. Conversely, ALKBH8, an enzyme responsible for tRNA modification, ensures translation accuracy and regulates oxidative stress by modifying the wobble position (e.g., mcm5U). Loss of ALKBH8 can result in neurological disorders and dysregulated oxidative stress responses. In a stress environment, accurate protein folding and translation are crucial. Although CCT and ALKBH8 operate at different molecular levels—CCT ensuring proper protein folding and ALKBH8 ensuring translation fidelity—they converge on the broader roles of stress management and protein quality control. Deficiencies in tRNA modification, such as those caused by ALKBH8 dysfunction, could increase the burden of misfolded proteins, thereby heightening the reliance on the CCT complex for correction. Thus, despite the moderate *Jaccard index*, the inferred connection suggests a meaningful biological interplay between these components.

**Figure 3.**
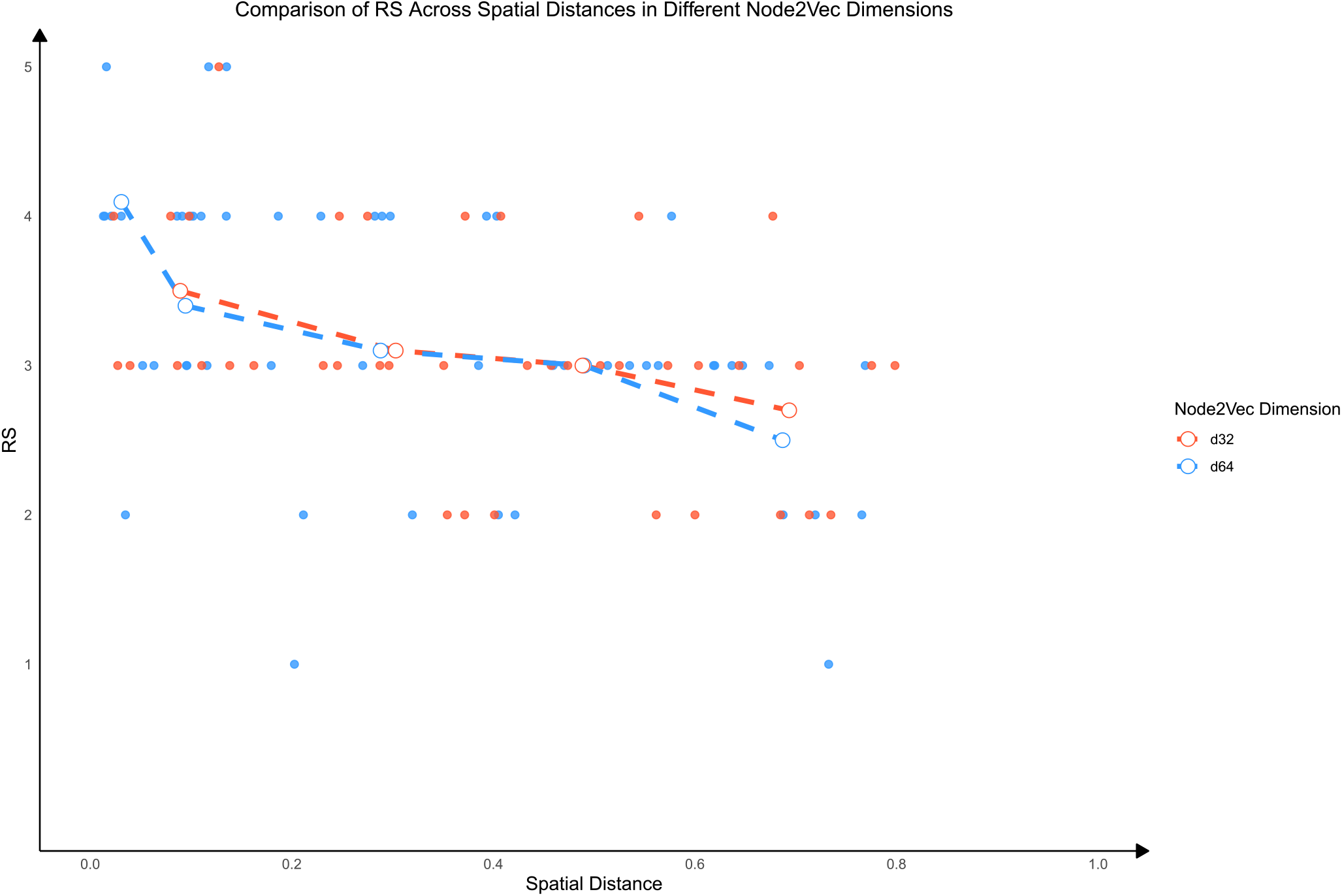
Linear correlation between reliability scores and node2vec distances in both 64- and 32-dimensional spaces, highlighting the superior performance of the 64-dimensional embeddings, particularly for pairs with distances below 0.1.

### Identifying Novel Biological Associations through Node2vec-Based Distances

Compared to traditional methods like the *Jaccard index*, which resemble differential gene analysis, node2vec results tend to have lower confidence scores based on literature surveys. However, they hold greater research value, as node2vec-derived spatial proximity often correlates with phenotypic synergy. Additionally, comparisons between the *Jaccard index* and node2vec spatial distances reveal that the 64-dimensional embeddings (d64) exhibit more extreme values (top 5%), indicating higher gene-level reliability. Furthermore, d64 scores consistently surpass d32 in terms of reliability (Figure 4), underscoring its superior performance in capturing meaningful biological relationships. we specifically focused on capturing those from the top 5% of the d64 distances. To further elucidate the relationships among terms, we developed a minimum spanning tree (MST, Supplementary Material 2) (Eisner, 1997) and an interactive web interface.

**Figure 4.**
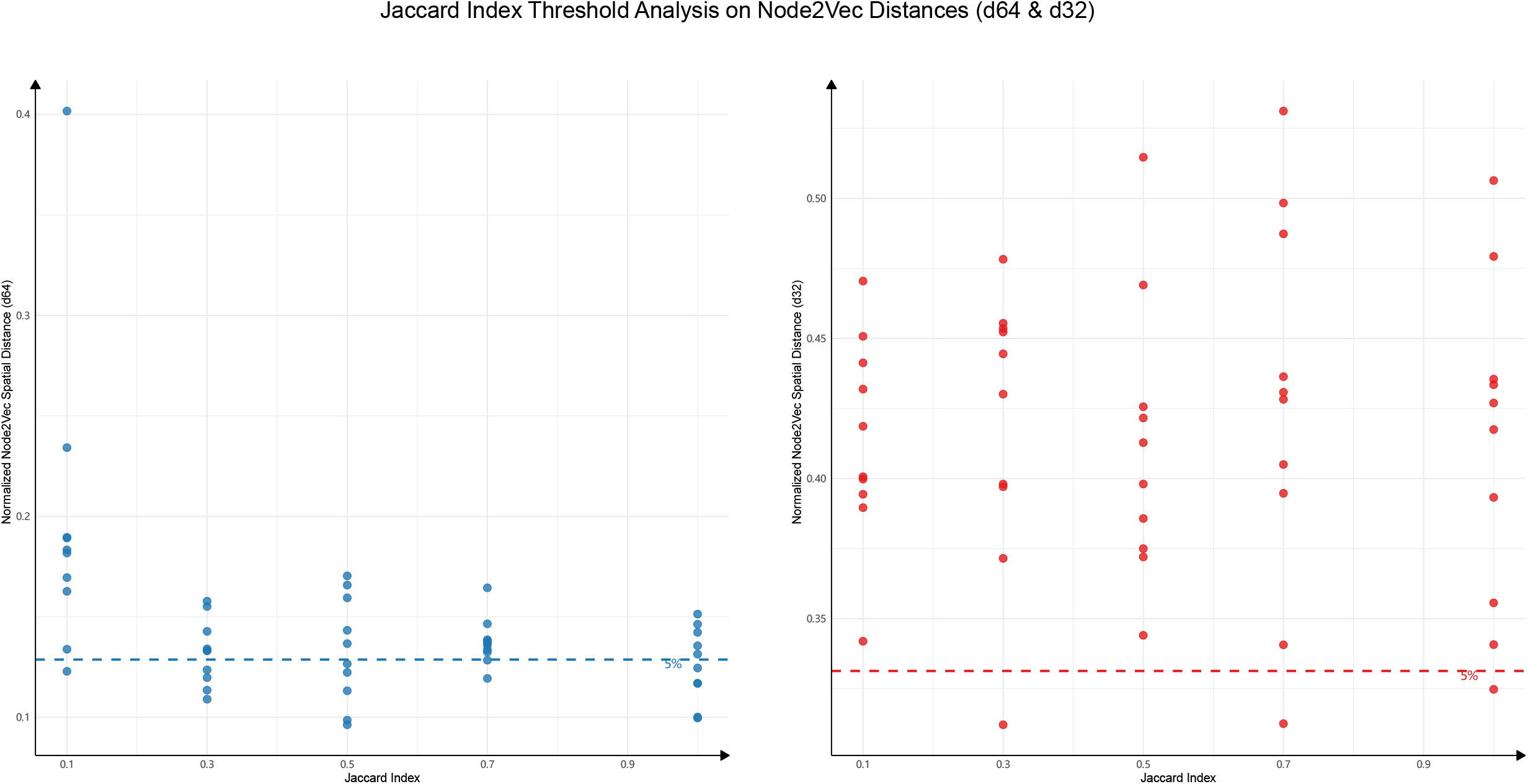
Correlation plot between the Jaccard index and both 64-dimensional (left) and 32-dimensional (right) node2vec distances, with dashed lines marking the top 5% : d64 = 0.13 and d32 = 0.33. Despite a relatively weak overall correlation, the plot supports the observation that smaller spatial distances in the 64-dimensional space are more likely to indicate biologically credible associations.

Our analysis reveals that a low Jaccard index does not invariably indicate a lack of functional or biological relevance, as several key genes across two distinct terms occupy crucial roles and upstream positions within the pathway. For instance, we observed a *Jaccard index* of merely 0.083 between the GNF2_PRDX2 gene set from the GNF2 expression atlas and the HP_ANISOCYTOSIS gene set. Despite the minimal overlap, the node2vec spatial distance between these terms ranks within the top 5%, hinting at a potential biological connection. This link may be driven by oxidative stress mechanisms, as PRDX2, an antioxidant enzyme, plays a pivotal role in regulating cellular redox balance. Oxidative stress is a key factor in various hematological disorders, and its modulation could influence red blood cell (RBC) morphology, specifically in conditions like anisocytosis, which is characterized by variations in RBC size. Altered PRDX2 expression has been associated with hematological conditions such as anemia, which frequently presents with anisocytosis. We hypothesize that PRDX2 may regulate RBC morphology by mitigating oxidative damage, thereby establishing a mechanistic link to anisocytosis. This potential connection between PRDX2 and RBC size variability could pave the way for further research into its role in hematological disorders, particularly those involving oxidative stress (Sadowska-Bartosz & Bartosz, 2023; Stresing *et al*., 2013).

Both KEGG_MEDICUS_REFERENCE_TYROSINE_DEGRADATION and WP_TYROSINE_METABOLISM_AND_RELATED_DISORDERS are pathways associated with tyrosine metabolism and degradation. The former emphasizes the overall metabolic process, while the latter focuses on the pathological mechanisms resulting from metabolic disorders. Although these pathways share an identical gene set, their spatial distance in the node2vec analysis ranks within the top 5%. This suggests a tight biological network connection between these pathways and hints at potential clinical phenotypic commonalities.

In contrast, despite having a Jaccard index of 1, KEGG_MEDICUS_REFERENCE_TRANSPORT_OF_CALCIUM and GOMF_GLUTAMATE_GATED_CALCIUM_ION_CHANNEL_ACTIVITY exhibit a noticeably larger spatial distance in the node2vec analysis and possess a RS that is four points lower than the others. The difference between these two terms lies in their perspective on the same activity; however, the former encompasses three fewer genes (*GRIA1, GRIA3, GRIK1*) than the latter. This highlights node2vec’s ability to distinguish between pathways based on their functional roles and pathological relevance, rather than being solely driven by gene set overlap. The KEGG pathway describes the broad physiological process of calcium ion transport across membranes, involving various cell types and tissues. In comparison, the GO molecular function pathway is more specific, focusing on the activity of glutamate-gated calcium ion channels within the nervous system, a more narrowly defined molecular mechanism. Consequently, node2vec separates these pathways in the biological network, reflecting differences in their functional and contextual roles despite their identical gene sets. This demonstrates node2vec’s capacity to capture topological distinctions in the network, offering a more nuanced understanding of functional relationships beyond simple gene overlap.

### Cross-Species Discovery Through Node2Vec Embedding Reveals Shared Biological Insights

Due to ethical constraints prohibiting functional perturbation experiments within human subjects, alongside the limitations of cell line experiments that diverge from the true *in vivo* environment, a substantial amount of research is conducted within murine models. The central aim of these animal experiments is to bridge murine data with human disease and physiological mechanisms, rendering cross-species association studies indispensable.

We extended our analysis to include cross-species validation, utilizing node2vec unsupervised embeddings at 64 dimensions. By embedding both human and mouse gene datasets into a shared vector space, we aimed to identify biologically relevant patterns that are conserved across species. Specifically, we converted mouse genes to their homologous human counterparts and embedded the mice collections based on the pre-trained human model. Following this, we searched for the nearest human term-related nodes corresponding to the embedded terms in mouse. This approach allowed us to discover and strengthen our findings across species (Supplementary Material 1).

The observed connection between mice_TABULA_MURIS_SENIS_LUNG_B_CELL_AGEING and human_CHARAFE_BREAST_CANCER_LUMINAL_VS_MESENCHYMAL_UP suggests potential links between ageing lung B cells and distinct cellular states in breast cancer(Almanzar *et al*., 2020). The mesenchymal transition in cancer is frequently associated with immune responses and cellular ageing, which might explain this correlation. Ageing B cells, particularly in tissues like the lung, may undergo significant functional changes, such as a reduction in antigen-specific responses or a shift towards pro-inflammatory profiles. These changes are characteristic of immunosenescence, which is known to both impair immune surveillance and enhance a pro-tumorigenic environment. Chronic inflammation, often seen with ageing immune cells, creates a microenvironment conducive to cancer progression through the secretion of inflammatory cytokines and growth factors that support tumor cell proliferation, survival, and metastasis. This is especially relevant to the mesenchymal transition in cancer, as inflammatory signals are critical drivers of epithelial-mesenchymal transition (EMT), facilitating cancer cell invasion and dissemination.

## Discussion

In this study, we utilized node2vec, comparing to straight forward set-overlapping methods, to explore complex biological relationships within diverse datasets. The key strength of this approach lies in its ability to embed terms from various datasets into a shared vector space, enabling researchers to discover relationships between similar terms across different datasets. This provides a flexible tool for uncovering hidden patterns, particularly in contexts where data is heterogeneous and often incomplete.

However, one of the limitation of this approach is the reliance on pre-existing datasets, which introduces a level of uncertainty. Our term embeddings are drawn from various sources, each with varying levels of curation and confidence, and we have not assigned explicit weights to these terms yet. This results in a broad, unsupervised analysis where both input and output lack precise control. While this introduces potential noise and ambiguity, it also allows for serendipitous discoveries, as seen in some of the unexpected relationships we uncovered. These findings suggest that even in a loosely structured exploration, novel biological insights can emerge, particularly when the direct evidence is limited or the underlying mechanisms are not well understood.

The node2vec algorithm itself alleviates some of the concerns mentioned above. Its simplicity and ease of parameter tuning make it well-suited for exploratory analyses. Additionally, its unsupervised nature is particularly advantageous when working with heterogeneous and incomplete data, which is common in biological research. This flexibility allows us to capture potential relationships that more rigid, supervised methods might overlook.

Despite these strengths, it is important to acknowledge that our study primarily serves as a proof-of-concept. The current framework demonstrates the potential of node2vec in biological research, but more concrete case studies with well-curated datasets are necessary to fully exploit its capabilities. Additionally, while node2vec offers simplicity and control, future work should explore the use of more sophisticated models, such as graph neural networks, which may offer greater precision and sensitivity when applied to specific, focused research questions.

## Code Availability

The code utilized in this study is available on GitHub at git@github.com:YuHang-aw /MsigDB_mining.git. It includes all necessary scripts for data processing, model training, and evaluation. The repository is licensed under the MIT license, facilitating reproducibility and supporting further research

## Data Availability

A user-friendly Excel file detailing the relationships between term pairs is available, including values such as the *Jaccard index*, 64-dimensional distances (d64), and 32-dimensional distances (d32), scaled to approximately 300. The datasets used and analyzed during this study are available in the Supplemental Data accompanying this article.

## Notes

https://github.com/YuHang-aw/MsigDB_mining.git

### Competing Interest Statement

The authors have declared no competing interest.

### Summary of Updates

Figures revised Reorganization Supplemental files updated

